# A Novel Efficient L-Lysine Exporter Identified by Functional Metagenomics

**DOI:** 10.1101/2020.04.30.071142

**Authors:** Sailesh Malla, Eric van der Helm, Behrooz Darbani, Stefan Wieschalka, Jochen Forster, Irina Borodina, Morten O. A. Sommer

## Abstract

Lack of active export system often limits the industrial bio-based production processes accumulating the intracellular product and hence complexing the purification steps. L-lysine, an essential amino acid, is produced biologically in quantities exceeding two million tons per year; yet, L-lysine production is challenged by efficient export system at high titres during fermentation. To address this issue, new exporter candidates for efficient efflux of L-lysine are needed. Using metagenomic functional selection, we identified 58 genes encoded on 28 unique metagenomic fragments from cow gut microbiome library that improved L-lysine tolerance. These genes include a novel putative L-lysine transporter, belonging to a previously uncharacterized EamA superfamily. Characterization using *Xenopus oocyte* expression system as well as an *Escherichia coli* host demonstrates activity as a L-lysine transporter. This novel exporter improved L-lysine tolerance in *E. coli* by 40% and enhanced the specific productivity of L-lysine in an industrial *Corynebacterium glutamicum* strain by 12%. Our approach allows the sequence-independent discovery of novel exporters and can be deployed to increase titres and productivity of toxicity-limited bioprocesses.

## Introduction

The global chemical industry is transitioning from reliance majorly on petrochemical processes to more sustainable bio-based production. This development holds promise to improve the sustainability of the chemical industry while also reducing the overall production costs of chemical products. In order to establish a cost competitive bioprocess, titers of fermentations frequently exceed 100 g/L, which leads to a significant stress on the host organism. Indeed, a majority of industrial bioprocesses are limited in production due to several stresses resulting from high product titers.

One of the most significant bio-based chemical products is L-lysine. The global bio-based L-lysine production now exceeds 2.5 million tons per year, which is estimated to reach 3.0 million tons in 2022 corresponding to 5.6 billion USD of market value according to the current L-lysine market report (Elder, 2019). Industrial L-lysine bioprocesses entirely rely on *Corynebacterium glutamicum* and *Escherichia coli* production strains that achieve titers over 1.2 M (Becker and Wittmann, 2012; Lee and Kim, 2015). It was observed that the L-lysine export rate is inhibited by 50% upon exceeding the extracellular concentration of 400 mM compared to that at 80 mM (Kelle et al., 1996), indicating the substantial inhibition of the L-lysine-specific export in industrial fermentation. Indeed, several studies have demonstrated the benefit of incorporating active efflux systems to address intracellular product accumulation in biobased production (Borodina 2019; Hemberger et al., 2011; Malla et al., 2010). Hence, L-lysine export system is an obvious target to maintain the producer organism at high lysine concentration as well as easing the downstream process. Despite its importance, there are only two identified lysine specific exporters: i) LysE as a member of the lysine efflux permease (LysE; **2.A.75**) family (Bellmann et al., 2001; Vrljic et al., 1999); and, ii) lysine outward permease (LysO or YbjE in *E. coli*) (Pathania and Sardesai, 2015). Vrljic et al. (1996) achieved five folds higher lysine export rate upon overexpression of LysE in *C. glutamicum* (Vrljic et al., 1996). Similarly, Yasueda and Gunji have deployed this strategy for ten-fold improvement in L-lysine production by expressing a spontaneously mutated LysE from *C. glutamicum* in *Methylophilis methylotrophus* (Gunji and Yasueda, 2006). In addition to the rational engineering of the existing exporters, there is an urgent demand for new genetic building blocks to further improve L-lysine tolerance and production.

Functional metagenomic selection is an effective approach to discover novel genes and enzymes due to its ability to access the wide range of genetic elements present in a particular environmental niche (Forsberg et al., 2016; Munck et al., 2015; Sommer et al., 2010, 2009). Hence, we set out to use functional metagenomic selection to identify novel L-lysine transporter candidates from a cow fecal library, with the goal of improving industrial L-lysine production (Fig. 1A). Using C13-labeled L-lysine and mRNA expression of the screened transporter in *Xenopus oocyte*, the transporter candidate was confirmed as L-lysine exporter. Expression of the metagenomic derived L-lysine transporter improved titer and productivity in both Gram-positive and Gram-negative production strains.

**Figure 1.**
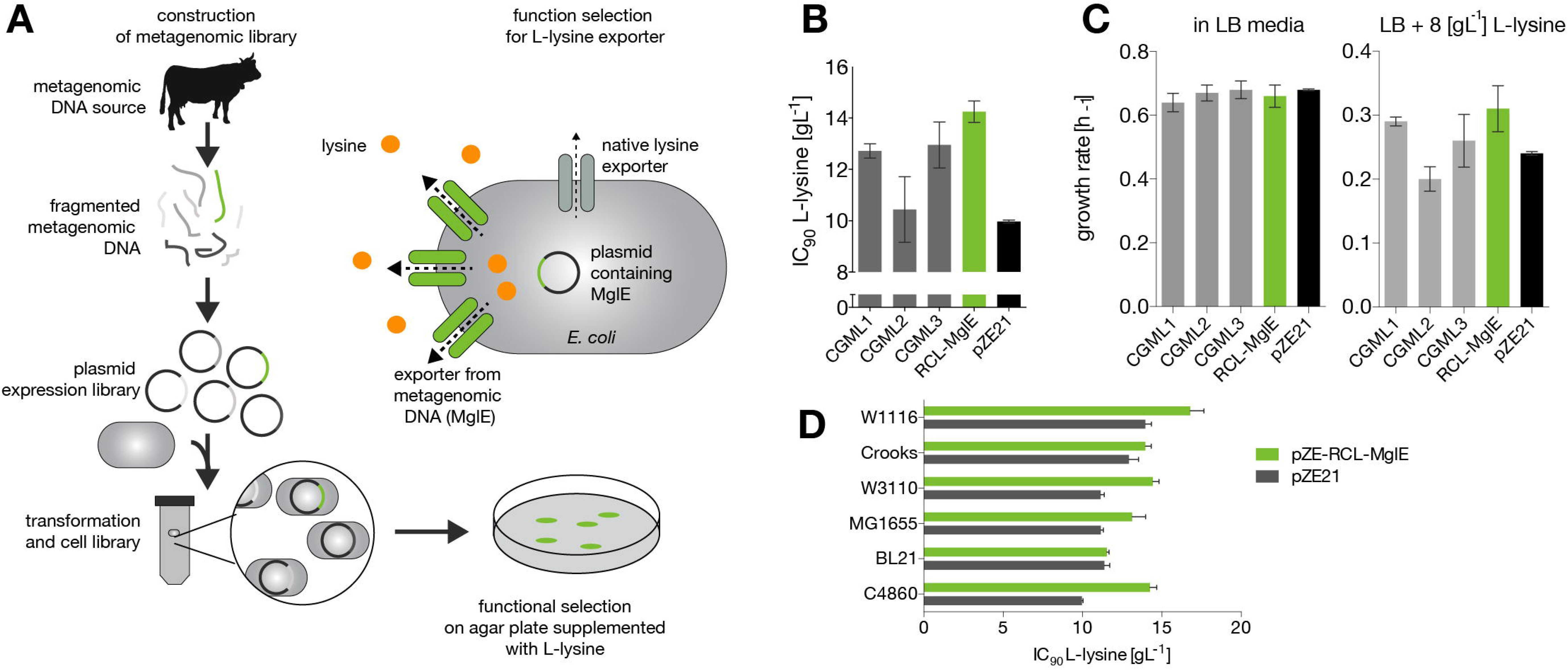
Functional metagenomic screening and Lysine tolerance of the screened metagenomics clones. A. Functional metagenomic screening for lysine transporters. Cow fecal metagenomic DNA library construction and its functional selection for L-lysine exporters. B. L-lysine IC90 values for the selected *E. coli* C4860 resistant clones harboring putative transporters. C. The growth rates of *E. coli* C4860 harboring pZE-RCL-MglE and pZE21 (control) in LB-Km and LB-Km supplemented with 8 g/L of L-lysine. D. Improved L-lysine tolerance (IC90 values) by expression of metagenomics insert carrying MglE transporter in industrially relevant *E. coli* strains.

## Materials and methods

### Bacterial strains, growth conditions and chemicals

All bacterial strains, vectors and plasmids used in this study are listed in Table 1. All oligonucleotide primers (synthesized by Integrated DNA Technologies, Inc.) used are presented in Table 2. *E. coli* strains were routinely cultured at 37°C in Luria-Bertani (LB) broth or on agar supplemented with kanamycin (35 μg mL^−1^) when necessary (hereafter referred as LB-km). For the L-lysine production in *E.coli* LB as well as M9 minimal media supplemented with 2 gm/L of yeast extract were used. *C. glutamicum* strains were cultured in modified CGXII medium (Keilhauer et al., 1993) supplemented with 5% sucrose, 1% BHI, 0.5 mM IPTG and kanamycin (when necessary), at 30°C and 250 RPM. Growth media were supplemented with 1.5% agar for plate assays.

**Table 1.**
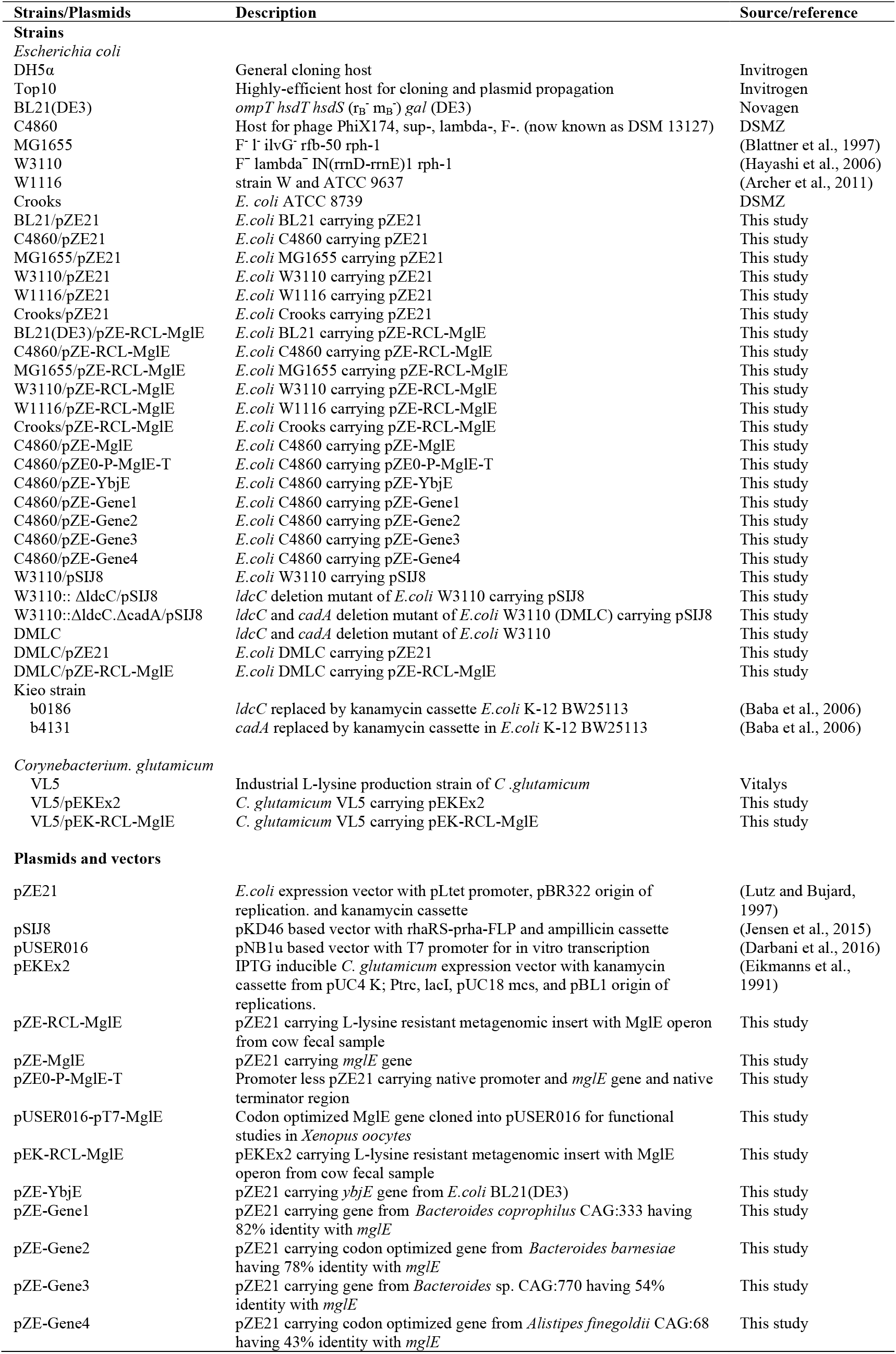
Bacterial strains and plasmids used in this study.

**Table 2.**
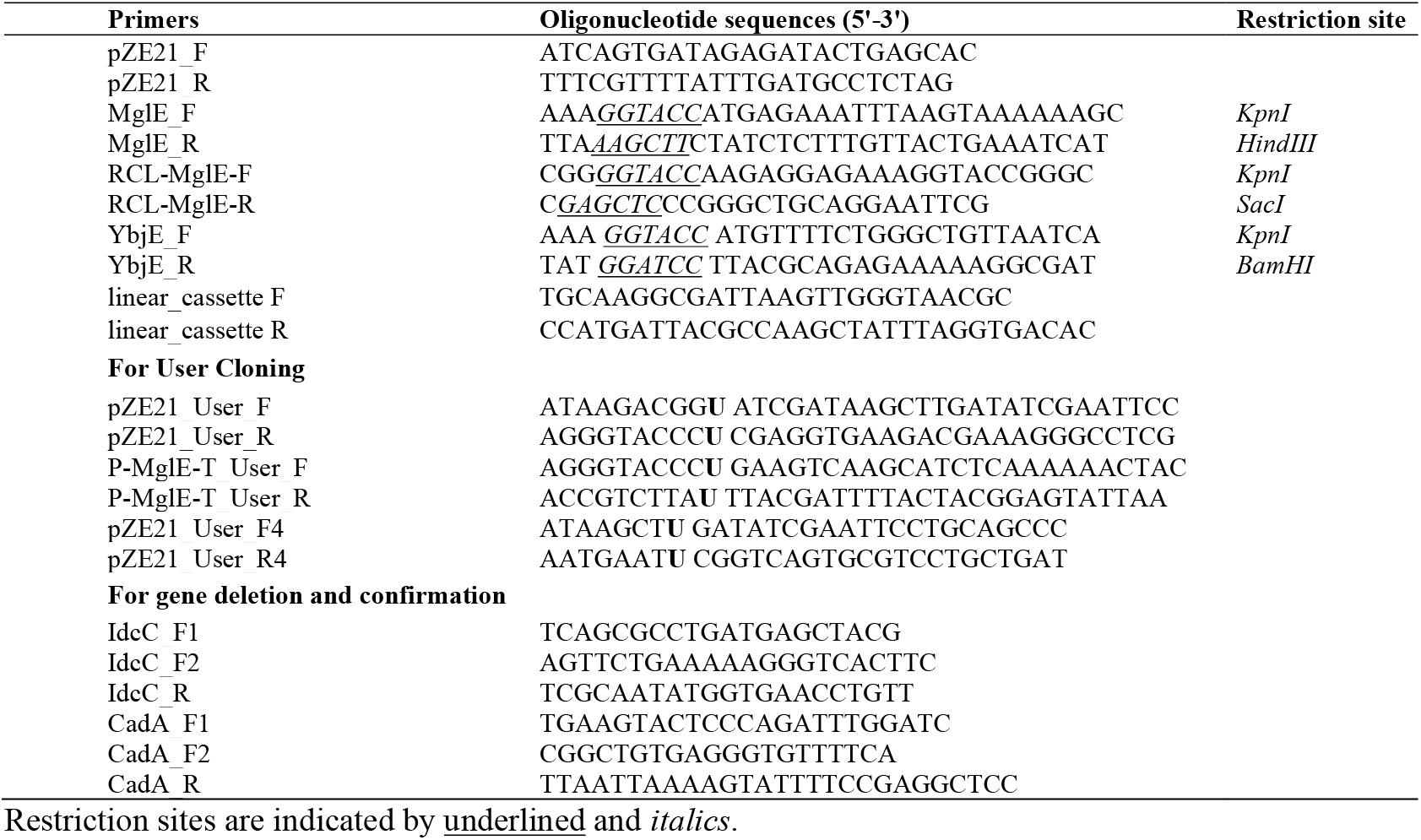
Oligonucleotides used in this study.

DNA manipulations were carried out following standard protocols (Sambrook and Russell, 2001). All chemicals were purchased from Sigma-Aldrich (St. Louis, MO, USA). Restriction enzymes and T4 DNA ligase were purchased from Fermentas (Denmark) and New England Biolabs (Hertfordshire, UK). DNA sequencing was performed using an automated DNA sequence analyzer.

### Metagenomic library construction and functional screening for L-lysine

A metagenomic expression library of total DNA extracted from a cow fecal sample was constructed as described previously (Sommer et al., 2009). Briefly, library construction involves i) the isolation of total DNA from 5 g of fecal material using the PowerMax Soil DNA Isolation Kit (Mobio Laboratories Inc.), ii) fragmentation of extracted DNA into pieces of an average size of 2 kb by sonication using a Covaris E210 (Massachusetts, USA) followed by end-repair using the End-It end repair kit (Epicentre), iii) blunt-end ligation into the pZE21 MCS1 expression vector with constitutive promoter pLteo-1 (Lutz and Bujard, 1997) at the *HincII* site using the Fast Link ligation kit (Epicentre), and iv) transformation of the ligated sample into electro-competent *E. coli* top10 cells by the standard method (One Shot® TOP10 Electropcomp™ Cells, Invitrogen).

After electroporation, cells were recovered in 1 ml of SOC medium for 1 h at 37°C, and the library was titered by plating out 1 μl, 0.1 μl and 0.01 μl of recovered cells onto LB-km plates. The insert size distribution was estimated by gel electrophoresis of the PCR products obtained by amplifying inserts using primers annealing to the vector backbone flanking the *Hinc*II site. The average insert size for the library was 1.7 kb. The total size of the metagenomics library was determined by multiplying the average insert size by the number of colony forming units (CFU) per ml (5×10^8^ bp). The remainder of the recovered cells were inoculated into 10 ml of LB-km liquid media and grown overnight; the library was frozen down in 15% glycerol and stored at −80°C.

The *E.coli* C4860 strain was used for the functional screening of L-lysine. First, a plasmid prep of the metagenomics library was carried out from the *E. coli* top10 cells harboring the library. Then, 400 ng of metagenomics plasmid DNA was transformed into electro-competent *E. coli* C4860 cells, and the library titer was determined as described above. On the basis of the determined library sizes and the titer of the library, 10^6^ cells (i.e., 100 μl of the library cells) were plated on LB-km agar supplemented with L-lysine at the selective concentration (14 g/L). Plates were incubated at 37°C, and the growth of colonies (likely lysine-tolerant clones) was assayed after 48-65 h of incubation.

The metagenomic inserts present in L-lysine-tolerant clones were Sanger sequenced using the pZE21_F and pZE21_R primer pair, which annealed to the vector backbone. The resulting raw sequencing chromatogram files were analyzed and functionally annotated using the deFUME web server (van der Helm et al., 2015): http://www.cbs.dtu.dk//services/deFUME/.

### Minimum inhibitory concentration (MIC) and IC90 determination

For MIC determination, the *E. coli* strains were cultured in LB liquid media at 37°C overnight, and then approximately 1×10^4^ cells were inoculated from the overnight cultures into LB liquid media and grown at 30°C and 300 RPM in 96-well microtiter plates containing 150 μl of medium per well. MICs were determined using a logarithmic L-lysine (or chemical) concentration gradient with two-fold serial dilutions. Endpoint absorbance measurements (A_600nm_) were taken with a plate reader (Synergy H1, BioTek) after 24 or 48 h of incubation and were background-subtracted. Growth inhibition of the *E. coli* strains was plotted against L-lysine (chemical) concentration with a polynomial interpolation between neighboring data points using R software (http://www.r-project.org). The percentage of inhibition was calculated using the formula: 1 − [A_600nm lysine (or chemical)_/A_600nm control_]. The inhibitory concentration was defined as the lowest concentration of the chemical that inhibited 90% of the growth of the strain tested (IC90).

### Growth experiments

Single colonies of the tolerant clone(s) harboring transporter homologues were grown overnight at 37°C with shaking in liquid LB-km medium. The OD values at 600 nm [(OD)_600nm_] were determined by 10-fold dilution. Then, the cultures were diluted to adjust the (OD)_600nm_ to 0.1, and 5 μl of each culture was inoculated in 150 μl fresh media containing L-lysine (0 and 8 g/L) and kanamycin in a 96-well micro-titer plate. The plate was incubated at 37°C for 24 h in an automated spectrophotometer (EL×808, BioTek) that recorded the (OD)_630nm_ at an interval of 60 min. The data were subsequently retrieved and analyzed to determine growth rates. The growth rate data are the average of triplicate experiments, with error bars representing the standard error of the mean (SEM).

### PCR and *E. coli* transformation

PCR was performed in a total volume of 50 μl under the following DNA amplification conditions: 95°C for 5 min, followed by 30 cycles of 95°C for 30 sec, 50–65°C for 30 sec, and 72°C for 1 min, and finally 72°C for 5 min. Electrocompetent or chemically competent *E. coli* cells were transformed with the ligation mixture using a standard protocol and plated onto LB-km agar plates for selection.

### *In silico* analysis

The amino acid sequence of MglE (**M**etagenomics **g**ene for **l**ysine **E**xport) was analyzed using Interpro (http://www.ebi.ac.uk/interpro/v), its 2D membrane topology was predicted using Phobius (Käll et al., 2007) and visualized using Protter (Omasits et al., 2014), and the phylogenetic tree was plotted using iTOL. The UniProtKB database (Consortium, 2014) was accessed on 2016-06-01 to query the closest homologs of MglE. Pfam version 29.0 (December 2015) (Finn et al., 2014) was used to extract the LysE family (Pfam id: PF01810) members. The Maximum Likelihood Phylogenetic tree was constructed using CLC Main Workbench 7 with the Neighbor Joining construction method.

### Transport assays in *Xenopus* oocytes

The *Xenopus laevis* oocytes were obtained from Ecocyte Bioscience (Germany). Oocytes were kept in Kulori buffer (pH 7.4) and at 18°C. A linear cassette (including T7 promoter, MglE as the gene of interest, and 3′UTR) was amplified from the plasmid pUSER016-T7-MglE using Phusion Hot Start polymerase (ThermoFisher Scientific) and the primers linear_cassette F and linear_cassette R. The linear cassette was used as template for *in-vitro* transcription. Capped cRNAs was synthesized using the mMESSAGE mMACHINE® T7 Transcription Kit (AM1344; ThermoFisher). The quality and quantity of the RNA were determined by Agilent 2100 Bioanalyzer. For microinjection of cRNA and 13C-labeled L-lysine, we used the RoboInject (Multi Channel Systems, Reutlingen, Germany) automatic injection system (Darbani et al., 2019). Injection needles with opening of 25 μm were used (Multi Channel Systems, MCS GmbH). For expression in oocytes, 25 ng of the *in-vitro* transcribed cRNA for the MglE transporter was microinjected into the oocytes 3 days prior to transport assays. To perform the export assay, 50 nl of the 13C-labeled L-lysine stock solution was injected into the oocytes to obtain estimated internal concentrations of 6 mM, assuming an after-injection dilution factor of 20 (Darbani et al., 2016). Following four washing steps, each batch of 20 oocytes was incubated for 180 min in 90 μl Kulori buffer at pH 5 or pH 7.4. After incubation, 70 μl of the medium was collected from each batch with intact oocytes and added onto 70 μl of 60% MeOH before LC-MS analysis. The oocytes were washed four times with Kulori buffer pH 7.4 and intracellular metabolites were extracted in 30% MeOH to analyzed on LC-MS.

### *E. coli* double deletion mutant construction

The chromosomal lysine decarboxylases *ldcC* and *cadA* in *E. coli* W3110 were successively knocked out by PCR targeting to create *E.coli* DMLC strain. The gene disruption process was carried out as described previously using an ampicillin-resistant pSIJ8 helper plasmid containing both the λ Red and FLP systems (Jensen et al., 2015). The primer pair IdcC_F2/IdcC_R (situated 64 bp away from the start/stop codon *ldcC* gene) was used to amplify the kanamycin cassette from the genomic DNA of the *ldcC* inframe knocked out Kieo strain b0186 whereas the primer pair CadA_F2/CadA_R (situated 56 bp away from the start/stop codon *cadA* gene) was used to amplify the kanamycin cassette from the genomic DNA of the *cadA* inframe knock-out Kieo strain b4131. These PCR products were used to delete *ldcC* and *cadA* in *E. coli* W3110 strain. The detail process of construction of *E. coli* DMLC is given in supplementary materials.

### Plasmid construction

#### Sub-cloning the metagenomic insert implicated in tolerance phenotypes

The recombinant plasmids pZE-MglE and pZE0-P-MglE-T were constructed as described below. The construction of recombinant plasmids was verified by both restriction mapping and direct nucleotide sequencing of the respective genes in the recombinant plasmids.

Using the oligonucleotide pair MglE_F/MglE_R and the pZE-RCL-MglE plasmid as template DNA, the exact *orf* of the putative carboxylate/amino acid transporter homologue (referred to as MglE) was amplified. The amplicon was purified with a Qiagen PCR purification kit and digested with the restriction enzymes *Kpn*I and *Hind*III (NEB, UK), followed by ligation using T4 DNA ligase (Fermentas, Denmark) into the corresponding restriction sites of the multiple cloning site (MCS) of the pZE21 vector to construct the recombinant plasmids pZE-MglE.

To construct the recombinant plasmid pZE0-P-MglE-T, USER cloning was applied. The pZE21-vector backbone without its promoter (pZE0) was amplified using the oligonucleotide pair pZE21_User_F/pZE21_User_R. The putative carboxylate/amino acid transporter homologue and the native promoter and terminator sequences (P-MglE-T fragment) were amplified from the pZE-RCL-MglE plasmid using the primer pair P-MglE-T_User_F/P-MglE-T_User_F. The amplified PCR products were ligated into the recombinant plasmid pZE0-P-MglE-T using the recommended standard USER cloning protocol (NEB, UK).

#### Codon optimized mglE plasmids construction for oocytes expression study

For functional analysis of MglE in *Xenopus* oocytes, we constructed recombinant plasmid pUSER016-pT7-MglE by USER cloning. The pUSER016 vector backbone was digested with *Pac*I for 18 h and the ends were further nicked with Nt.BbvCI for two hours. The codon optimized *mglE* gene (sequence provided in supplementary data) for *Xenopus laevis* was synthetized along with N-terminal linker sequence-GGCTTAA**U** and C-terminal linker sequences -**A**TTAAACC (in the complementary sequence, T is replaced by U for pairing with **A**) including uracil with compatible USER sites easing direct cloning into linearized and nicked pUSER016 vector by USER cloning (USER^®^, NEB), resulting in plasmid pUSER016-pT7-MglE.

#### Recombinant plasmids with E.coli lysine exporter and homologous genes of mglE

The lysine exporter, *ybjE*, (900 bp, Genbank accession no. CAQ31402) from *E.coli* BL21 (DE3) was amplified using oligonucleotides YbjE-F and YbjE-R. The PCR product of *ybjE* was cloned into pZE21 vector excised with *Kpn*I and *BamH*I restriction enzymes to construct pZE-YbjE expression plasmid.

Similarly, four *mglE* homologous genes; i) Gene1 (891 bp, 82% identity, Genbank accession no. CDC57518) from *Bacteroides coprophilus* CAG:333, ii) Gene2 (891 bp, 78% identity, Genbank accession no. WP_018711839) from *Bacteroides barnesiae*, iii) Gene3 (870 bp, 54% identity, Genbank accession no. CDC66277) from *Bacteroides sp.* CAG:770, iv) Gene4 (912 bp, 43% identity, Genbank accession no. CCZ76555) from *Alistipes finegoldii* CAG, were synthetized along with N-terminal linker sequence - AATTCAT**U** AAAGAGGAGAAAGGTACC and C-terminal linker sequence - GTCGACGGTATCG **A** TAAGCTT (in the complementary sequence, T is replaced by U for pairing with **A)** including uracil easing direct user cloning. These four synthetized gene fragments were cloned into the linear PCR fragment of pZE21 vector amplified by using oligonucleotides pZE21_User-F4 and pZE21_User-R4 (Table 2) to construct pZE-Gene1, pZE-Gene2, pZE-Gene3 and pZE-Gene4 recombinant plasmids, respectively.

### Construction of recombinant industrial *Corynebacterium glutamicum* strains

The full metagenomics insert derived from pZE-RCL-MglE plasmid was amplified using the primer pair RCL-MglE-F/RCL-MglE-R. The resulting PCR product was cloned into pEKEx2 vector, using the *Kpn*I and *Sac*I restriction sites, yielding expression plasmid pEK-RCL-MglE. The newly constructed expression plasmid was introduced into the industrial L-lysine producing strain *C. glutamicum* VL5 by electroporation and concomitant selection on kanamycin containing 2xTY-Agar. Successful construction of the new strain, named *C. glutamicum* VL5/pEK-RCL-MglE, was verified by re-isolation of the plasmid, followed by control digest and PCR on the length of the insert. The same procedure was performed to introduce the empty pEKEx2 expression vector into *C. glutamicum* VL5, yielding control strain *C. glutamicum* VL5/pEKEx2.

### Analysis of L-lysine in supernatant of bacterial culture

A calibration curve of an authentic standard was generated using concentrations ranging from 0.4 to 77 mg/L. The accurate mass of L-lysine from 20-fold diluted samples (supernatants containing L-lysine for quantification) was analyzed using an LC-MS Fusion (Thermo Fisher Scientific, USA) with positive electrospray ionization (ESI+). The final concentration was adjusted for the dilution factor. Bracketing calibration was used for the quantification of the external concentration. For the quantification of L-lysine, an LC-MS/MS, EVOQ (Bruker, Fremont USA) was used with multiple reaction monitoring (MRM) transition in positive ionisation mode (ESI+), with the quantified transition 147→84 (CE 10) and qualifier transition 147→130 (CE 7). The significance of the specific lysine production was calculated using a Student t-test.

## Results and Discussion

### Identification of metagenomic L-lysine tolerance genes

The gut microbiota from industrial livestock animals are conceivably enriched with the microbes capable to survive at higher L-lysine concentration which is supplied as food additives. A metagenomic library derived from cow feces was constructed and used to find candidate genes that could lead to a higher L-lysine tolerance in *E.coli* C4860 strain. The resulting library was screened for L-lysine tolerance by plating on LB agar plates supplemented with inhibitory concentrations of L-lysine (Fig. 1A) (materials and methods). Colonies appeared only on the plates with the metagenomic library but not on the control plates with *E.coli* C4860 harboring empty vector. Eighty L-lysine tolerant clones were selected for further analysis.

Metagenomic inserts present in those L-lysine tolerant clones were PCR amplified, sequenced and annotated using deFUME (van der Helm et al., 2015). In total, 28 unique clones containing 58 complete or partial open reading frames were identified in this analysis. The sequence analysis revealed that resistant clones contained *orf*s homologous to genes encoding L-lysine modification/degradation enzymes, membrane proteins, signaling proteins and hypothetical proteins. (Supplementary Table S1).

### Selection of transporter candidates

For the purpose of improving industrial production strains, novel efflux systems constitute more relevant genetic building blocks than those involved in degradation and modification, which would counteract the objective of L-lysine production. Accordingly, we focused our subsequent analysis on *orfs* encoding potential efflux systems. We identified six unique metagenomic inserts encoding putative membrane proteins, some of which are annotated in Genbank as hypothetical proteins (Supplementary Table S1). Four inserts were selected for further testing based on the absence of putative degradation enzymes flanking the putative transporters on the metagenomic insert.

We determined the L-lysine IC90 values for each of the four selected metagenomic inserts. The L-lysine IC90 values of the selected metagenomic clones ranged from 10.44 ± 1.277 g/L to 14.25 ± 0.415 g/L, corresponding to a more than 40% increase in the L-lysine IC90 for the clone harboring pZE-RCL-MglE compared to the empty vector control strain (Fig. 1B). All of these strains have similar growth profiles in LB medium, whereas the growth rates varied in the liquid LB supplemented with 8 g/L of L-lysine. At 8 g/L L-lysine supplementation, the growth rate of the highest tolerant metagenomic clone harboring pZE-RCL-MglE was 30% higher than that of the empty vector control (Fig. 1C).

### Lysine tolerance by MglE in *E. coli* strains

To test whether the pZE-RCL-MglE plasmid carrying the *mglE* gene could confer L-lysine tolerance to industrially relevant *E. coli* strains, we introduced the plasmid into various *E. coli* strains; BL21 (DE3), MG1655, Crooks, W1116, and W3110 and IC90 values of L-lysine were determined for all of the constructed *E.coli* strains. Interestingly, the IC90 values of L-lysine for all of these *E. coli* strains were increased upon introduction of the pZE-RCL-MglE plasmid (Fig. 1D). Of particular interest, the L-lysine IC90 was increased by 30% in W3110, which is a widely used background strain for L-lysine production (Imaizumi et al., 2005). These data demonstrate that the mechanism of L-lysine tolerance mediated by pZE-RCL-MglE is effective across the industrially relevant *E. coli* strains and accordingly should be generally applicable to industrial *E. coli-*based L-lysine fermentations.

### Protein sequence analysis of the metagenomic insert carrying MglE

Sequence analysis of the 1.6 kb PCR amplicon from pZE-RCL-MglE metagenomics insert contained an open reading frame *mglE* flanked by native promoter and terminator sequences (Fig. 2A). The MglE protein consists of 297 amino acids, and its closest homolog is a hypothetical protein from *Bacteroides coprophilus* (Genbank accession no. WP_008140691) with 82% identity at the amino acid level. The MglE protein is a member of RhaT/EamA-like transporter family of the drug/metabolite transporter (DMT; **2.A.7**) superfamily. The MglE contains two copies of the EamA domain, which is found in transporters belonging to the EamA family, at 9–143 aa (pfam (Finn et al., 2014) e-value: 3e^−11^) and 152–292 aa (pfam e-value 9.6e^−12^). The members of the EamA family are diverse, and most of their functions are unknown (Franke et al., 2003). Nevertheless, a few proteins belonging to EamA family are well characterized as exporters such as PecM exports a pigmented compound indigoidine in *Erwinia chrysanthemi* (Rouanet and Nasser, 2001) and YdeD export metabolites of the cysteine pathway in *E. coli* (Daßler et al., 2000). The predicted two-dimensional topology of the MglE transporter possesses six cytoplasmic domains, five periplasmic domains, and ten transmembrane domains, with both the N- and C-terminals in the cytoplasmic region (Fig. 2B).

**Figure 2.**
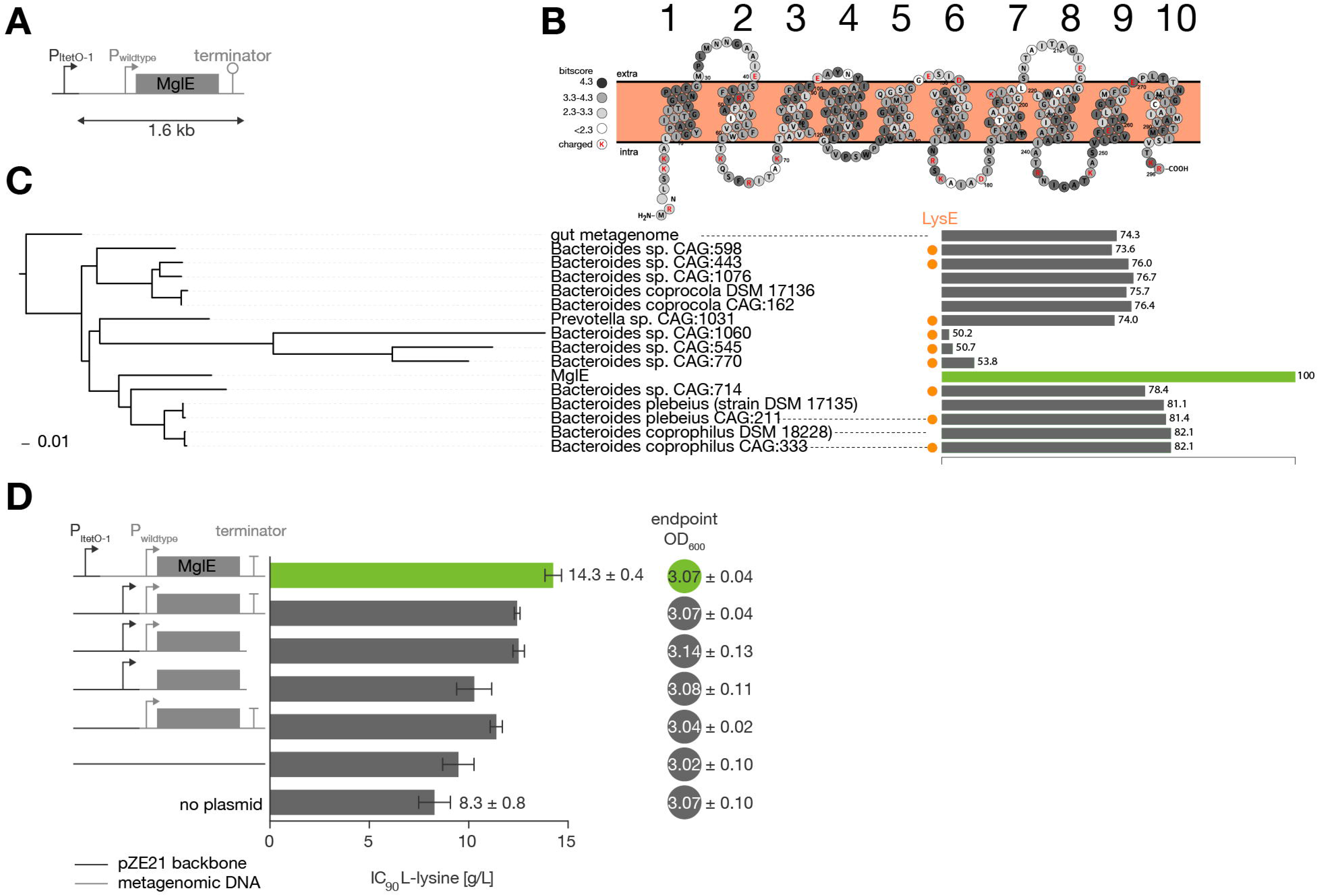
*In silico* analysis of the metagenomic insert contained in pZE-RCL-MglE plasmid and L-lysine tolerance of its subclones. A. The metagenomic insert present in the pZE-RCL-MglE plasmid is shown. P: promoter region, MglE: membrane bound transporter protein, and T: terminator sequence. B. The membrane topology of the MglE transporter protein showing 6 cytoplasmic regions, 5 periplasmic regions, and 10 transmembrane α-helices. The 15 BLASTp hits with a >50% sequence identity were aligned to MglE and the conservation (bitscore) for each residues was calculated using the Biopython package (Cock et al., 2009). C. A Maximum Likelihood phylogenetic protein tree containing all the significant (sequence identity >50%) BLASTp hits in the UniProtKB database against MglE. The percentage identity of the 15 hits is shown as horizontal bars. The presence of a LysE family member in the genomes is shown with an orange dot. The highest homology with MglE is found in *Bacteroides coprophilus* CAG:333, with 82% sequence identity. D. The strategy for subcloning the MglE transporter with or without its native promoter and the resultant L-lysine tolerance phenotypes for the *E. coli* C4860 strain are shown. The biomass of the strains (at 24 h) cultured in LB media was also shown in terms of OD in closed circles.

Phylogenetic analysis shows that the closest MglE homologs (>50% sequence identity) are present in the phylum Bacteroidetes and are annotated as uncharacterized proteins (Fig. 2C). Members of the LysE family (pfam id: PF01810) are mainly (>5 species per phyla) found in the phyla: Proteobacteria, Bacteroidetes, Firmicutes, Actinobacteria. The full list of phyla that contain species with a LysE domain are Proteobacteria: 102, Bacteroidetes: 73, Firmicutes: 45, Actinobacteria: 28, Chloroflexi: 1, Cyanobacteria: 3, Spirochaetes: 1, Verrucomicrobia: 1 and Gemmatimonadetes: 1. Interestingly, nine genomes containing the MglE homolog also contain additional LysE family (pfam id: PF01810, TC code: **2.A.75**) genes (Fig. 2C orange dot). MglE represents a novel L-lysine transporter that could not have been identified by sequence analysis or homology to known L-lysine transporters.

### Effect of regions in the metagenomics insert on L-lysine tolerance

To investigate the effects of promoter region of the metagenomic clone on lysine tolerance, the recombinant plasmids pZE-MglE (*mglE* cloned into pZE21), and pZE0-P-MglE-T (MglE along with its native promoter and terminator cloned into pZE21 without the pLteo-1 vector promoter) were subcloned from pZE-RCL-MglE plasmid. The constructed plasmids were transformed into *E. coli* C4860, and IC90 values of L-lysine were determined (Fig. 2D). Although these recombinant C4860 strains grew in the same extent in LB media, they showed different L-lysine tolerance. We observed that the IC90 for the strain harboring the pZE0-P-MglE-T plasmid was higher than that of the strain harboring the pZE-MglE plasmid, indicating that the native promoter resulted in higher lysine tolerance than the vector promoter. However, the synergistic effect of these two promoters was better still (i.e., the IC90 values of the pZE-RCL-MglE plasmid was higher than those of the strains harboring pZE-MglE or the pZE0-P-MglE-T plasmids). Hence, the lysine tolerance of the *E. coli* C4860 strain was the highest upon harboring the full metagenomic insert (i.e., pZE-RCL-MglE).

### MglE exports L-lysine when expressed in *Xenopus* oocytes

For functional analysis, MglE was expressed in *Xenopus* oocytes and the export of C13-labeled L-lysine was measured. The water-injected oocytes were used as negative control. L-lysine export assay was performed by injecting C13-labeled L-lysine to obtain a final cytosolic concentration of 6 mM in the oocytes. The oocytes were incubated in Kulori buffers with two different pH (pH 5 and 7.4) for 3 h, and L-lysine concentrations were measured in the buffer and within the oocytes. MglE-expressing oocytes resulted in 6.6- and 8.5-fold higher extracellular concentrations of C13-labeled L-lysine and natural C12 L-lysine respectively than control oocytes, when buffer with pH 5 was used (Fig. 3Ai). The intracellular concentrations of C13-labeled and natural C12 L-lysine were correspondingly 2.1- and 1.4-fold lower than in control oocytes (Fig. 3Ai). On the other hand, there was no significant change in L-lysine quantities when oocytes were incubated at pH 7.4 (Fig. 3Aii). This indicates a proton-gradient dependent export mechanism for MglE transporter. L-lysine resistance (uptake) experiments in *E. coli* also indicated a proton-dependent uptake for L-lysine because L-lysine toxicity was higher at pH 8.5 and 10 than at pH 4.5 and 7 (data not shown).

**Figure. 3.**
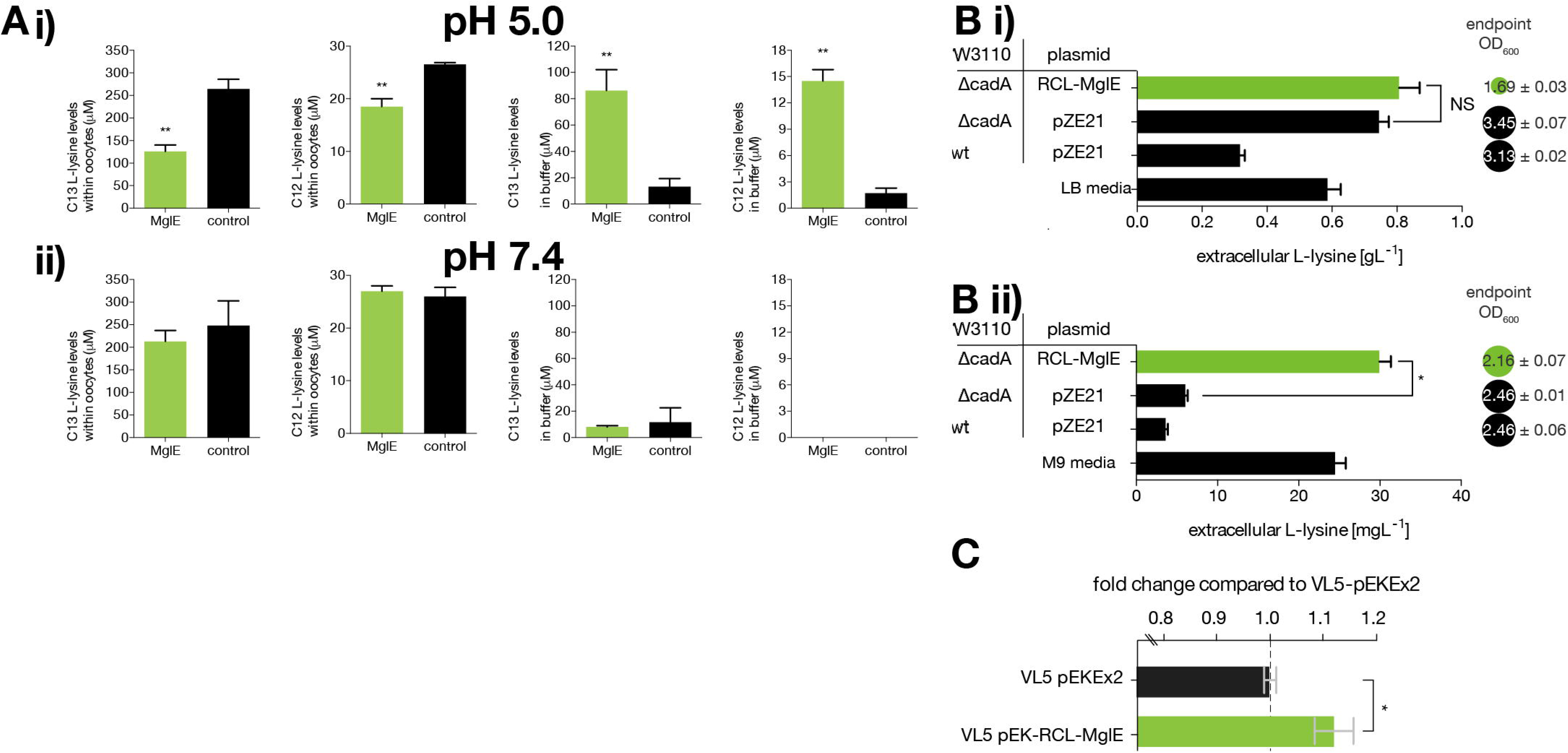
Functional characterization of MglE transporter activity in *Xenopus* oocytes, *E.coli* and *C. glutamicum* A. L-lysine export assay of MglE transporter in *Xenopus* oocytes in Kulori buffer i) at pH 5 and ii) at pH 7.4. The bars represent lysine contents in the oocytes or the buffer (means, ± Std., 3-4 biological replicates each involving 20 oocytes). Significant changes are in comparison with the controls marked by asterisks (** *p* < 0.01; Fishers one-way ANOVA). B. The extracellular L-lysine concentration of reference *E. coli* W3110 and its isogenic mutant strain DMLC along with the cell OD_600nm_ values confirming the exporter activity of the MglE protein in LB media and ii) in M9 minimal media supplemented with yeast extract (* p<0.0001, t-test). C. Fold enhancement of specific productivity of L-lysine by MglE protein in the industrial *C. glutamicum* VL5 strain compared to the empty vector control. Error bars are s.e.m, * denotes p = 0.023, n=4).

### MglE assists L-lysine export in *E.coli*

To analyze the function of MglE for L-lysine efflux, *E. coli* DMLC strain was constructed by knocking out two lysine decarboxylase genes, constitutive (*ldcC*) and acid-inducible (*cadA*). These deletions should prevent L-lysine degradation, thereby leading to increased L-lysine production (Kikuchi et al., 1997). Then, the pZE-RCL-MglE plasmid and pZE21 empty vector were transformed into *E. coli* DMLC, yielding *E. coli* DMLC/pZE-RCL-MglE and *E. coli* DMLC/pZE21, respectively. Subsequently, these strains along with *E. coli* W3110/pZE21, were cultured in LB media as well as M9 minimal media supplemented with yeast extract. The bacterial growth and extracellular L-lysine concentrations were measured after 24 h. We found that in LB media the absolute extracellular L-lysine titer was increased from 744 mg/L to 806 mg/L in presence of the metagenomic insert consisting MglE operon i.e. 8.3% higher L-lysine production in the *E. coli* DMLC/pZE-RCL-MglE strain as compared to *E. coli* DMLC/pZE21 in LB. However, the biomass of the strain expressing MglE was decreased by about 50% as compared to the control strain (Fig. 3Bi). Furthermore, the L-lysine production was analysed in minimal media supplementing yeast extract. In minimal media, the extracellular L-lysine accumulation was increased from 6 mg/L to 30 mg/L (p<0.0001, t-test) whereas there was 12% reduced in biomass upon expressing the MglE metagenomic fragment (Fig. 3Bii). The difference in extracellular L-lysine concentration among the strain expressing MglE and the control strain is prominent in minimal media (p<0.0001*, t-test) than in LB media (p=0.1997, t-test). Hence, the above results demonstrate that MglE assists exporting L-lysine from *E. coli*.

The reduced biomass might be due to the combined effect of expression of membrane protein in high copy number and active export of L-lysine by MglE which could have resulted in less available energy for the cell to build biomass. It is worth noting that a big proportion of total cellular energy-demand is for the transport machinery which has accordingly been under evolutionary selection towards a higher energetic efficiency (Darbani et al., 2018).

### Improvement of L-lysine productivity in industrial *C. glutamicum* strain

Expression of a functionally active gene(s) from one strain to another often requires special optimizations. For the practical application in industrial L-lysine bioprocesses, retaining the activity of the discovered exporter into the production host, *C. glutamicum*, is very essential. To test whether the MglE can boost the L-lysine bioprocesses, we cloned and expressed it in an industrially used *C. glutamicum* L-lysine production strain.

The full metagenomic insert carrying the MglE along with its native promoter and terminator was amplified from the pZE-RCL-MglE plasmid and cloned into pEKEx2 expression vector to construct pEK-RCL-MglE expression plasmid. The constructed plasmid was introduced into *C. glutamicum* VL5 strain (Materials and methods), a producer strain currently used for industrial L-lysine production. Using sucrose as a carbon source, L-lysine production from the constructed *C. glutamicum* recombinant strain was analyzed at 30 h. Unlike in *E.coli* strains, expression of the full metagenomics insert harboring MglE did not have any growth inhibition effect as the vector control and the recombinant strains both have nearly equal biomass. Notably, upon incorporation of the MglE operon into *C. glutamicum*, the specific L-lysine productivity was improved by 12 ± 0.07% relative to the empty vector control (Fig. 3C). By expressing the operon in this highly efficient producer strain, we showed not only the functionality of the MglE in a Gram-positive species but also the benefit of the efflux system on an already highly optimized producer strain.

### Comparison of L-lysine tolerance conferred by MglE and its homologous genes

The YbjE transporter from *E. coli* is one of the functionally characterized lysine-specific exporters. Based upon the homology search, four genes in the NCBI database having different homologies (ranging from 43% to 82% in amino acid level) with MglE were also selected and gene synthetized. To find out the lysine tolerance of MglE as compared to that of YbjE and the four selected MglE homologous proteins (with 82, 78, 54 and 43% amino acid sequence identity), recombinant expression plasmids pZE-YbjE, pZE-Gene1, pZE-Gene2, pZE-Gene3 and pZE-Gene4 were constructed. The constructed plasmids, pZE-MglE and pZE21 vector control were introduced into *E.coli* C4860 and *E.coli* DMLC strains and determined the IC90 of L-lysine for the constructed recombinant strains as described in early experiments.

Interestingly, we found that the expression of MglE provided similar L-lysine tolerance in *E.coli* C4860 strain as that of YbjE whereas in *E.coli* DMLC strain, MglE displayed even higher tolerance than YbjE. On the other hand, all of the four MglE homologs showed lower lysine tolerance than MglE in both strains (Fig. 4).

**Figure. 4.**
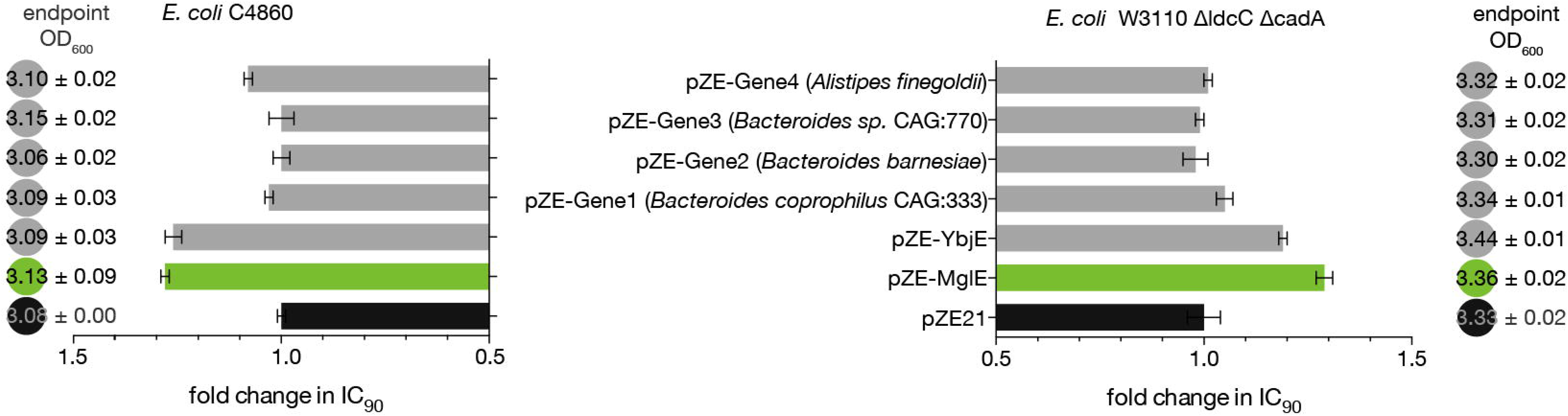
Comparison of lysine tolerance (presented in relative IC90 values) displayed by MglE, YbjE (Lysine specific exporter from *E.coli*) and four synthetic MglE homologs. The biomass of the strains (at 24h) cultured in LB media was also shown in closed circles.

## Conclusion

We deployed functional metagenomic selections to identify novel genetic building blocks encoding product efflux systems. Through applying this approach to the problem of L-lysine toxicity at high concentration, we discovered MglE, which is confirmed as a novel type of L-lysine exporter that belongs to the EamA superfamily. We demonstrate the benefits of MglE in both Gram-negative (*E. coli*) and Gram-positive (*C. glutamicum*) strains and show that MglE expression improves the productivity of an industrial L-lysine production strain. If incorporated into industrial-scale bioprocesses, this discovery has the potential to enhance L-lysine production by about 12%, representing an increased profit on the order of 200-500 million USD per year. Our approach of using functional metagenomics to identify novel transporters is independent of prior knowledge of transporter gene sequences and can be generally applied to most of the toxic compounds/chemicals production by fermentation.

### Protein accession numbers

The sequence information of full metagenomics insert harboring *mglE* is deposited in the NCBI GenBank database with the accession number KU708839.

## Acknowledgements

The research leading to these results was funded by the Novo Nordisk Foundation (Grant Agreement no. NNF10CC1016517), and the European Union Seventh Framework Programme (FP7-KBBE-2013-7-single-stage) under grant agreement no. 613745, Promys. This work was further supported by the European Union Seventh Framework Programme-ITN (FP7/2012/317058 to E.v.d.H). Stefan Wieschalka acknowledges funding from Højteknologifonden. IB acknowledges the financial support the European Research Council under the European Union’s Horizon 2020 research and innovation programme (YEAST-TRANS project, Grant Agreement no. 757384). We would like to thank Vitalys I/S for providing the VL5 strain for lab-scale testing, Christian Munk for helping with R-script to calculate IC90 values.

## Competing financial interests

The authors declare no competing financial interests.

## Author contributions

M.O.A.S. and S.M. conceived the study; S.M. designed and performed *E.coli* experiments, B.D and I.B. conceived the experiments on *Xenopus oocytes* and B.D. performed experiments. S.W. performed *C. glutamicum* experiments, E.v.d.H performed *in silico* analysis. M.O.A.S., I.B., and J.F. led the research teams. M.O.A.S. and S.M. drafted the manuscript and the other authors contributed to the writing.

## Notes

### Competing Interest Statement

The authors have declared no competing interest.

