## Supplementary file 1 for "A Novel Efficient L-Lysine Exporter Identified by Functional Metagenomics"

**Supplementary Table S1**. List of orf and homologous protein in lysine resistant clones

| Metagenomic plasmid clone | **Occurance** | **Most similar protein GenBank accession no** | **Organism (accession no. of similar protein)** | **e-value** | **Query coverage %** | **% Identity (a.a.)** |
| --- | --- | --- | --- | --- | --- | --- |
| pZE-CGML_1_ | 5 | hypothetical protein CD30_04750  peptidase C15  multidrug transporter  hypothetical protein | *Lysinibacillus massiliensis* CCUG 49529 (KGR91620)  *Bacillus* sp. FF3 (WP_042473155)  *Bacillus isronensis* (WP_008407819)  *Lysinibacillus manganicus* (WP_036181882) | 1.1E-09  5.3E-40  2.9E-16  2.5E-12 | 6.07  47.7  16.5  92.3 | 60  68  72.5  59 |
| pZE-CGML_2_ | 2 | oligopeptide/dipeptide ABC transporter ATPase subunit  peptide ABC transporter substrate-binding protein | *Subdoligranulum* sp. CAG:314 (WP_022466037)  *Clostridium* sp. M2/40 (WP_044038993) | 2.1E-94  1.6E-37 | 37.8  24.8 | 75.6  71.6 |
| pZE-CGML_3_ | 8 | hypothetical protein | *Tannerella* sp. CAG:51 (WP_022391164) | 1.9E-13 | 47.8 | 30.8 |
| pZE-CGML_4_  Renamed as  pZE-RCL-MglE | 8 | hypothetical protein (with Carboxylate/Amino Acid/Amine Transporter domain) named as MglE | *Bacteroides coprophilus* CAG:333 (WP_022277040) | 1E-142 | 99 | 82 |
| pZE-CGML_5_ | 2 | transporter  acetyltransferase (GNAT) family  tRNA (guanine-N(1)-)-methyltransferase | *Marvinbryantia formatexigens* (WP_006864112)  *Faecalibacterium* sp. CAG:82 (WP_022256375)  *Clostridiales bacterium* oral taxon 876 (WP_021655073) | 8.5E-26  1.5E-61  5.1E-29 | 30.1  81.9  30.5 | 52.1  68.2  68.5 |
| pZE-CGML_6_ | 2 | hypothetical protein  hypothetical protein  MFS transporter | *[Ruminococcus] gnavus* (WP_024854345)  *Gilliamella apicola* (WP_034885103)  *Bifidobacterium adolescentis* (WP_039775353) | 3.9E-23  3.9E-09  2.3E-43 | 62.1  96.1  32.9 | 35  38.2  55.3 |
| pZE-CGML_7_ | 6 | GNAT family acetyltransferase  hypothetical protein  hypothetical protein | *[Clostridium] hathewayi* (WP_025530003)  *Clostridium paraputrificum* (WP_027098406)  *Paenibacillus dendritiformis* (WP_006677989) | 2.1E-23  5.4E-51  4.1E-12 | 28.4  106  38.2 | 81.8  47.6  30.5 |
| pZE-CGML_8_ | 4 | GCN5 family acetyltransferase  Fic family protein  hypothetical protein | Bryobacter aggregatus (WP_031495868)  Prevotella bryantii (WP_027452747)  Odoribacter splanchnicus CAG:14 (WP_022160830) | 3.00E-20  9.00E-46  2.60E-62 | 28  44  47 | 67  57  55.9 |
| pZE-CGML_9_ | 4 | Protein of unknown function (DUF1653)  GCN5 family acetyltransferase  flavoprotein family protein  cytidylate kinase | *[Clostridium] termitidis* (WP_004628977)  *Shewanella* sp. ECSMB14102 (WP_039033060)  *Clostridium* sp. CAG:1013 (WP_016405150)  *Ruminococcus* sp. CAG:488 (WP_022081453) | 1E-23  2.1E-35  4.7E-27  1.2E-51 | 87.3  71.8  17.7  64 | 66.7  53.2  63.5  58 |
| pZE-CGML_10_ | 12 | ornithine cyclodeaminase | *Lachnospiraceae bacterium* 6_1_37FAA (WP_008977234) | 2.6E-117 | 64.2 | 66.3 |
| pZE-CGML_11_ | 1 | alanine racemase  alanine dehydrogenase | *Caldicellulosiruptor bescii* (WP_015908285)  *Sedimentibacter* sp. B4 (WP_019228209) | 2.1E-40  8.5E-88 | 36.7  48.6 | 51.4  69.8 |
| pZE-CGML_12_ | 5 | tRNA (guanine-N1)-methyltransferase  tyrosyl-tRNA synthetase | *[Clostridium] josui* (WP_024834621)  *Butyricicoccus pullicaecorum* (WP_016146822) | 9.2E-103  3.6E-147 | 96.5  57 | 59.9  84.6 |
| pZE-CGML_13_ | 1 | prolyl-tRNA synthetase | *Ruminococcus* sp. CAG:488 (WP_022079938) | 1.7E-101 | 38.1 | 79.2 |
| pZE-CGML_14_ | 1 | 16S rRNA methyltransferase  methionyl-tRNA formyltransferase | *Thermosediminibacter oceani* (WP_013275831)  *Ruminococcus* sp. CAG:724 (WP_021997445) | 4.8E-35  2.2E-77 | 36.2  78 | 40.4  51.2 |
| pZE-CGML_15_ | 1 | oxidoreductase  hypothetical protein | *Bacteroidales bacterium* ph8 (WP_019129248)  *Alistipes senegalensis* (WP_019151155) | 8.5E-105  5E-84 | 71.9  43.3 | 75.1  53 |
| pZE-CGML_16_ | 1 | Clp protease ClpC | *Bacteroides faecichinchillae* (WP_025074169) | 4.9E-107 | 23.3 | 80.7 |
| pZE-CGML_17_ | 1 | pyridoxamine 5'-phosphate oxidase family protein | *Peptoclostridium difficile* (WP_021414695) | 9.4E-34 | 54.2 | 61.5 |
| pZE-CGML_18_ | 1 | ribonucleoside-triphosphate reductase  hypothetical protein | *Lachnoclostridium phytofermentans* (WP_029503774)  *Clostridium* sp. CAG:277 (WP_022464576) | 4.2E-144  1.3E-145 | 32.6  34.5 | 85.8  82.4 |
| pZE-CGML_19_ | 1 | hypothetical protein, partial  nitroreductase family protein | *Prevotella* sp. oral taxon 473 (WP_044062834)  *Faecalibacterium* sp. CAG:74 (WP_022445666) | 1.2E-34  4.2E-49 | 83  89.1 | 37.4  51.9 |
| pZE-CGML_20_ | 1 | MULTISPECIES: N-acetylgalactosamine-6-sulfatase  hypothetical protein | *Bacteroides* (WP_008761730)  *Catenibacterium mitsuokai* (WP_006505171) | 3.5E-09  6.2E-19 | 9.84  49.2 | 56.9  65.6 |
| pZE-CGML_21_ | 1 | hypothetical protein  UDP-N-acetylglucosamine 2-epimerase | *Parabacteroides* sp. CAG:409 (WP_022455774)  *Parabacteroides goldsteinii* (WP_007658584) | 1.5E-24  4.2E-134 | 32.1  95.3 | 40.1  56.3 |
| pZE-CGML_22_ | 1 | NAD-dependent dehydratase  hypothetical protein, partial | *Lachnospiraceae bacterium* AB2028 (WP_027435031)  *Clostridium* sp. ATCC 29733 (WP_021660244) | 1.8E-33  9.3E-23 | 27.7  95.5 | 64.4  51.2 |
| pZE-CGML_23_ | 1 | nicotinate phosphoribosyltransferase  hypothetical protein | *Bacteroides* sp. CAG:1060 (WP_021843767)  *Coprobacter fastidiosus* (WP_022600833) | 1.1E-61  1.9E-19 | 44.3  28 | 58.2  43.8 |
| pZE-CGML_24_ | 3 | site-specific recombinase, phage integrase family | *Clostridium* sp. ATCC BAA-442 (WP_021632979) | 1.2E-09 | 8.05 | 78.1 |
| pZE-CGML_25_ | 1 | glycerophosphoryl diester phosphodiesterase | *Ruminococcus* sp. CAG:177 (WP_022188323) | 8.8E-40 | 27.7 | 71 |
| pZE-CGML_26_ | 2 | hypothetical protein | *Bacteroides fragilis* (WP_032544072) | 0.000015 | 56.8 | 41.2 |
| pZE-CGML_27_ | 1 | conserved hypothetical integral membrane protein  hypothetical protein  hypothetical protein | *Butyrivibrio* sp. CAG:318 (WP_022261702)  *Tenacibaculum maritimum* (WP_024742374)  *Prevotella* sp. CAG:1058 (WP_021852775) | 8.4E-61  3.8E-11  3.6E-85 | 68.4  30  60.6 | 59  44.6  53.1 |

**Construction of *E.coli* DMLC**

The chromosomal lysine decarboxylases *ldcC* and *cadA* in *E. coli* W3110 were successively knocked out by PCR targeting. The gene disruption process was carried out using an ampicillin-resistant pSIJ8 helper plasmid containing both the λ Red and FLP systems (Jensen et al., 2015). In the pSIJ8 plasmid, genes for λ Red recombinase (γ, β, *exo*), which enhance the recombination rate, were cloned under the control of the arabinose promoter, while FLP recombinase was cloned under the control of the rhamnose promoter to eliminate the resistance cassette.

*ldcC* disruption: The primer pair IdcC_F2/IdcC_R (situated 64 bp away from the start/stop codon *ldcC* gene) was used to amplify the kanamycin cassette from the genomic DNA of the *ldcC* inframe knocked out Kieo strain b0186. First, the pSIJ8 plasmid was transformed into the *E. coli* W3110 strain using chemically competent cells. The transformed cells were then selected on LB-amp solid medium and incubated at 30°C. The *E. coli* W3110/pSIJ8 strain was grown in LB-amp at 30°C to an OD_600nm_ of 0.3; then, 1 mM arabinose was added to induce the λ Red system. Next, electrocompetent cells were prepared following a standard protocol. Approximately 200 ng (5 µl) of the purified PCR product was added to 50 µl of competent cells in an ice-cold cuvette, and the transformation process was carried out as described previously. The transformed cells were plated on LB-km, amp solid medium. To confirm the replacement of *ldcC* by the kanamycin cassette, colony PCR was performed using the primer pair IdcC_F1/IdcC_R, which yielded a 1.6 kb band corresponding to the km cassette, thereby indicating the replacement of the *ldcC* gene (2.1 kb). Then, the colony confirmed to harbor a replacement of *ldcC* by a km cassette was grown in 1 ml of LB-km-amp at 30°C for 3–4 h. The cells were collected by centrifugation and resuspended in 100 µl of sterilized water; then, 10 µl of the resuspended cells was inoculated into 1 ml of LB-amp containing 50 mM of rhamnose and incubated at 30ºC for 4–6 h. The cells were spread onto an LB-amp plate and incubated at 30°C overnight. To confirm the removal of the km cassette, colony PCR was performed using the primer pair IdcC_F1/IdcC_R, which yielded a ~0.25 kb rather than a 1.6 kb band, confirming the successful deletion of the resistance marker. For further confirmation, the colonies were streaked on LB-amp and LB-km plates. The deletion mutant grew only on LB-amp but not on LB-km. The strain was named *E. coli* W3110::Δ*ldcC*/pSIJ8.

*cadA* disruption: Similarly, the primer pair CadA_F2/CadA_R (situated 56 bp away from the start/stop codon *cadA* gene) was used to amplify the kanamycin cassette from the genomic DNA of the *cadA* inframe knock-out Kieo strain b4131. The purified PCR product was transformed into *E. coli* W3110::Δ*ldcC*/pSIJ8, and a similar protocol was followed for the confirmation of *cadA* deletion. To confirm the replacement of *cadA* by the km cassette as well as the removal of the km cassette in the double deletion mutant, the primer pair CadA_F1/CadA_R was used for colony PCR.

To remove the helper plasmid pSIJ8 from the double deletion mutant, *E. coli* W3110::Δ*ldcC.*Δ*cadA*/pSIJ8 was grown in LB medium at 37°C for 4–6 h; then, 50 µl of 1000- and 10,000-fold diluted cells were spread onto an LB plate. The plates were incubated at 37°C overnight, and the next morning, the colonies were streaked on LB and LB-amp plates. The pSIJ8 lost strains grew on LB but not on LB-amp. This *ldcC* and *cadA* deletion strain was named *E. coli* DMLC.

**Nucleotide sequences:**

1) MglE operon where *mglE* gene is shown in blue font

>AAATATTATCTATTTTTATATTCCTCAGACGAAAAAGAGCCTCCTTTTCACAGAATCCCATTATTTTAGGCTATGAAAAATCACATTTTCCCTCAGGGGCTAATGGCAATTACAAGTACGTGGGAACACCGGCTACCGACATGTTTGCCAACCGCGAACCTGTCATGCTGGAATGATGTGAACCAACGGTCGACGCCCGAGGCATCAAGCCGTAGGGGCTGAAGTCAAGCATCTCAAAAAACTACGGAGCTGTCGTTCTGGCAGAGCGTAATTGTTTAAATAAAATTCAAACAGCTTTTAAGAATAGTGGGAGAATATCAATATCTTTGCACTCCGATAATGTGATTGTTGATTTTATGAGAAATTTAAGTAAAAAAGCCCTTATTGGTTATCCGGCCGGAATAATCACGGGCATAACATACGGTCTCAACCCGCTGTTCGGAATGCCGCTGATGAATAACGGTGCGGCCATAGAGTCAATACTGTTCTTCCGCTATGCGTTTGCAGTGGTAATCCTGGGCGTGTTCCTTTGGCTCACAAAACAGAGTTTCAGGATTACAGCAAAGCAGACCGCTGTGCTGCTGGTGCTGGGCTTGCTCTACACTGCAAGCAGTCTATTCCTCTTCGAGGCATACAACTATATCGCCTCGGGATTGGCTACCACGCTAATATTCCTTTATCCTGTACTGGTGGCTGTTATTATGGTCTTTCTGGGTGTCGTGCCCTCGTGGCCCGTATGGCTTGCCATAGCAGCTATATTCGGCGGCGTACTTATTATGACACAAGGCAGCGGCGGCGAGTCGATAGATCCCGTCGGTGTCCTGTTGTCGCTTGGCTCGGCACTGGTCTATGCGCTCTTTATCGTCATCATCAACCGCAGCAAGGCTATTGCCGACATATCAAACTCACTGCTCACCTTCTATGCCTTGACAGTAGGCGCAATAGTATTTCTTGGGAAGATTGCCCTCTCCAACACGGCTATCACGGCAGGCATCGAGGGTGGGGCGGCATGGCTCAATCTTATAGGCCTGGCGTTGCTGCCGACAATCGTCTCCACCGCATCTCTTGCCATAGCTACACGCAACATAGGTGCAACAAAAGCGTCAGTGCTGGGCGTGTTTGAGCCCATAACGGCAATCCTTGTCGGAACAGTGATGTTCGGCGAGCCACTCACCACAAACATCATGCTCGGCATCTGCATCGCAATTATGGCAGTCACATTTATGATTTCAGTAACAAAGAGATAGAAACAGCGGAATAAACATATACAAGCGGCTGAGGAATAATCGCCTTCTCCATAATTTGTTTCTTTTTTCCATCGTCCCTTTCAATTATATTGGTTAGGAACAATTGTGTGTATGCCCGCATCAGAGATATCTACAGAGCATTTGTCCGAGGCTTACAAATAAATTTTGTAGTTGTTGGACAAATGCTTTGCCAACTTAATCCCATCTTGTTTACGCCTTATGTATAGCTAACCCCCATCTTACTTACAATCTATAAGGCTAAATGCACATTATTCCACAATGGGGAGGTGTTTTTCTTCCTCTTTTTCTCTTTATATTGCAATATTATTTTAATACTCCGTAGTAAAATCGTAAATAA

2) The codon optimized version of MglE transporter for oocyte experiment.

>ATGAGAAATcttAGTAAGAAAGCCCTTATTGGTTATcctGCCGGAattATCactGGCatcACATACGGTCTCAACcctCTGTTCGGAATGcctCTGATGAATAACGGTgctGCCattGAAtccatcCTGTTCTTCCGCTATgccTTTGCAGTGGTTATCCTGGGCGTGTTCCTTTGGCTCACAAAGCAGAGTTTCAGGATTACAGCAAAGCAGACCGCTGTGCTGCTGGTGCTGGGCTTGCTCTACACTGCAAGCAGTcttTTCCTCTTCGAGGCATACAACTATATCGCCtctGGATTGGCTACCaccctgattTTCCTTTATCCTGTCCTGGTGGCTGTTATTATGGTTTTTCTGGGTGTGGTGccatccTGGCCCGTTTGGCTTGCCATAGCAGCTattTTCGGCGGCGTTCTTATTATGACACAGGGCAGCGGCGGCGAGtctatcGATccaGTCGGTGTGCTGTTGtccCTTGGCtctGCACTGGTTTATgctCTCTTTATCGTTATCATCAACCGCAGCAAGGCTATTGCCGACattTCAAACtctCTGCTCACCTTCTATGCCTTGACAGTTGGCGCAatcGTTTTTCTTggaAAGATTGCCCTCTCCAACactGCTATCaccGCAGGCATCGAAGGTggagcaGCATGGCTCAATCTTattGGCCTGgctTTGCTGcctACAATCGTCTCCACCGCATCTCTTGCCATAGCTACACGCAACattGGTGCAACAAAAgcttctGTGCTGGGCGTGTTTGAACCCatcactGCAATCCTTGTCGGAACAGTGATGTTCGGCGAGCCACTCACCACAAACATCATGCTCGGCATCTGCATCGCAATTATGGCAGTCACATTTATGATTTCAGTGACAAAGAGAtaa

3) MglE homologies

i) YbjE gene

>ATGTTTTCTGGGCTGTTAATCATTCTGGTTCCCCTGATTGTGGGTTACCTCATTCCGCTTCGCCAACAAGCTGCGTTAAAAGTTATTAATCAGCTATTAAGCTGGATGGTTTACCTTATTCTCTTTTTTATGGGTATCAGTCTGGCGTTTCTCGATAACCTCGCCAGTAACCTGTTGGCGATTCTGCATTATTCTGCCGTCAGTATTACCGTTATTTTACTGTGTAATATTGCCGCCCTGATGTGGCTGGAGCGAGGCCTGCCGTGGCGCAACCACCATCAGCAAGAAAAACTCCCGTCGCGTATTGCGATGGCGCTGGAGTCGCTAAAACTGTGCGGCGTAGTAGTGATTGGTTTTGCCATTGGTCTAAGTGGACTGGCTTTCTTACAACACGCGACCGAAGCCAGTGAATACACGTTAATTTTGCTACTTTTCCTCGTTGGTATTCAGTTGCGCAATAATGGCATGACCTTAAAGCAGATTGTCCTTAATCGCCGGGGAATGATTGTCGCCGTGGTGGTGGTTGTCAGTTCATTAATTGGTGGTTTAATTAACGCCTTTATTCTTGATCTCCCCATCAATACCGCGCTGGCAATGGCCTCCGGTTTCGGCTGGTATTCTCTTTCCGGTATTTTATTGACCGAATCTTTTGGTCCGGTAATCGGGAGCGCGGCGTTTTTTAATGATCTGGCCCGTGAACTGATTGCTATTATGTTGATCCCTGGGCTGATTCGCCGCAGCCGCTCTACTGCACTGGGCTTATGCGGTGCCACATCAATGGATTTCACCCTGCCCGTTCTTCAACGTACTGGCGGGCTGGATATGGTCCCGGCGGCAATTGTTCACGGTTTTATTCTTAGCCTGTTAGTGCCGATCCTCATCGCCTTTTTCTCTGCGTAA

ii) Gene 1 (82% identity with MglE)

>ATGAAAAAATTAAGTAAAAATGCCATTATCGGTTATCCTGCCGGAATTATTACCGGTATTACTTACGGATTAAATCCTCTATTTGCGGTTCCACTGATGAAGAATGGTGCCGCCACTGAGTCCATACTTTTTTTCCGCTATGCCTTTGCAGTACTTCTTCTGGGTCTGTTTTTAATGCTCCGCAAACAAAGTTTCCGGATCTCAGGAAAGCAAATAGGTGTATTGCTTGCACTCGGAGTTCTTTATACTTCAAGCAGCGTTTTTCTTTTCGAGGCCTATGAATACATCGCATCGGGACTGGCCACAACCCTGGTCTTTTTATATCCCGTCCTTGTGGCTATCATCATGGTATTCCTGAAAGTCGTCCCCTCCTGGCCTGTCTGGCTTGCCATTGCGGCAACCTTCGGAGGAGTGCTGGTAATGACACAAAGCGACGGTACACAGACCATCAACCCTGTCGGCGTATTGCTTTCAATTGCCTCGGCACTGGTATACGCACTCTTCATTGTCATCATCAACCGTAGCAAAGCCATTGCCGGCATCTCCAATTCATTGCTTACCTTCTATGCGCTTACGGTGGGAGCAATCGTGTTTCTCGGAAAAATCATCTGTTCCGACACCGCCATAACCGCCGGAATCACAACCGGAGCAGACTGGCTTAATCTCGTAGGACTTGCCCTGTTACCGACCATTGTTTCCACGGCTACTCTTGCTATTGCTTCCCGAAACATCGGAGCAACCAAGGCATCAGTCCTCGGCGTCTTCGAACCCATCACGGCCATTGTTGTCGGCACACTTATGTTCGGCGAACCGCTTACAACAAATATCATTGTCGGAATCTCAATCGCCATGGTAGCCGTCACATTCATGATTACCGTGACCAAGCGATAA

iii) Gene 2 (78% identity with MglE

>ATGAAAAAACTGAGCAAAAACGCCCTGATTGGTTATCCGGCAGGTATTATTACCGGTATTACCTATGGTCTGAATCCGCTGTTTGCAGTTCCGCTGATGAAAAATGGTGGTGCAATTGAAAGCATCCTGTTCTTTCGTTATTTCTTTGCCGTTCTGATGCTGGCAGCATTTCTGGTTATTCGTAAACAGCGTTTTCGCATTAGCGGTAAACAGGCAGGCATTCTGCTGGTTCTGGGTCTGCTGTATACCGCAAGCAGCCTGTTTCTGTTTGAAGCCTATCATTATATTGCAAGCGGTCTGGCAACCACCCTGGTTTTTCTGTATCCGGTTCTGGTTGCCATTATTATGGTGTTTCTGAAAGTTGTTCCGAGCTGGCCTGTTTGGCTGGCAATTGCAGCAACCTTTGGTGGTGTTCTGATTATGACCCAGGGTGATGGTGCACAGGCACTGAACCCGCTGGGTGTTCTGCTGAGCCTGGGTAGCGCACTGGTTTATGCACTGTTTATTGTGATTATCAATCGCAGCAAAGCCATTGCCAGCATTAGCAATAGCCTGCTGACCTGTTGTACCCTGGTTGTTGGCACCTTTGTTTTTCTGGGTAAAATTCTGTATAGCGGCACCGAACTGACCAGCTGTATTACCACCGGCACCGATTGGTTTAATCTGCTGGGACTGGCACTGCTGCCGACCGTTGTTAGCACCGCAACCCTGGCAGTTGCAAGCCGTAATATTGGTGCAACCAAAGCAAGCGTTCTGGGTGTTTTTGAACCGATTACCGCAATTCTGGTTGGCACCCTGATGTTTGGTGAACCGCTGACCGCAAATATTGTTATTGGCATTGTGATTGCCATTGTCGCCGTTACCTTTATGATTACCGTTACCAAACGTTAG

iv) Gene 3 (54% identity with MglE)

>ATGGTATGGGGATTCGCTGCAGGAATCATTACCGGTGTCACTTATGGTCTCAATCCTCTTTTTGCGAAACCTCTTCTGCAGATGGGTGTTTCGGTAGATTCAATGCTTTCTTTCCGTTATCTTGTAGCGTGTCTGATTCTCGGTGTCTGGCTGCTTCTGAGGAAGGAAACTTTCAAGGTCAACAGGCCTCAGTTTTTCCGTTTGTGTATTCTGGGTGTCCTTTTTGCGCTGAGCAGCATGTTGCTGTTCCTTTCCTATAAGTATATCCCTGCCGGTCTTGCCACGACCATTGTTTTCCTTTATCCGGTTCTTGTGGCTTTCATCATGGTTTTCCTCAAGGTTTATCCTACGTGGCAGGTCTGGGTGTCTATTTTTCTGACTTTCGTCGGAGTGGTGATTCTCAGCCGCCCTTCCGGCAATGTTTCCCTCAATGCCGTGGGGCTTCTTCTTGCCGGCGGTTCGGCATTGGCCTATGCCCTTTATCTTGTAGTGGTGAACCGGAGCCGTCGTCTCCGTACCGTTTCCAATCATCTCTTGACTTTCTATGCGCTTCTTATCGGTTCCTTGGTGTTCCTTCTGCATAATCTCATTGGCGGCGGCGGTCTTATGACCGGTATTCACGGCTGGTATTGCTGGTTCAATATCATCGGTCTTGCCATATTCCCGACTCTTGTCTCGCTGCTGACTCTTGCCATCGCGACTCGTATCATCGGTGCGACCCGCACCTCTGTTCTCGGTGTGTTCGAGCCTGTGACCGCGATTGCCGTCGGCACAATCTTCTTCGGAGAGTCGCTTACTGTCAATGTTATTGTGGGCGTTGTAATTACTCTGGTTGCCGTGACGTTCATGGTCCTGACCGGCAAGAAGTAG

v) Gene 4 (43% identity with MglE)

>ATGGCGAACGCTAAGTCGCGCGGCTGCGTGCTGGGTGCCGTGGCCGCCGCCAGTTACGGATTGAATCCGCTCTTCACGTTGCCCCTCTACGAGGCGGGCATGGGAGTGGATTCGGTGCTTTTCTACCGTTATCTGCTGGCTGCGGCGATGCTGGGCGCGTTGATGCTCGTCCGGCGGCAGTCGTTCGCCGTGCGCCGCCGCGACCTTGTGCCGCTGGCGGTCATGGGACTGCTCTTTTCTTTCTCGTCGCTCTTCCTGTTCGAGAGCTACAACCACATGGACGCGGGCATCGCTTCGACCATCCTGTTCCTCTATCCCGTGCTGGTCGCGGTCATCATGGCCGTCGGTTTCCACGAGAAGGTGAGCCGGATCACCATGCTCTCGATCCTGCTGGCCTTCACCGGCATCGCCATGCTCTACAAAGGCGGCGGTGAGCCGCTCTCCTTTCTCGGCGTCGCGCTGGTTTTTCTCTCGTCGCTCTGCTATGCGGTCTACATCGTGGGCGTCAACCGCTCGTCGCTGCGCGGCCTGCCGACCGAGAAACTCACCTTTTACGCTCTGCTGTTCGGCCTGAGCGTCTATGTCGTGCGCCTGCGCTTCTGCGCCGACCTGCAGGCGATCCCCACGCCCGGACTCTGGATCAACGCCGTCTCGCTGGCGCTCTTTCCGACGATCGTGTCGCTGGTGACGATGGCCGCCGCCATCCGCGCCATCGGCTCTACGCCCACGGCGATCCTCGGTGCGCTGGAACCCCTTACGGCCCTCTTTTTCGGCGTCGTGGTCTTCGGCGAACGGCTCACGCTGCGCATCGTGCTGGGCGTGGTGCTGATTTTGGTTGCCGTGACGCTCATCATCGCCGGTCGGTCACTGCACATTCCTTTCCCGCACCTGCGGTTCCGCCGGGCGCGGTAG
